# Microbiota in milk from healthy and mastitis cows varies greatly in diversity, species richness and composition, as revealed by PacBio sequencing

**DOI:** 10.1101/2020.08.13.249524

**Authors:** Teng Ma, Lingling Shen, Qiannan Wen, Ruirui lv, Qiangchuan Hou, Lai Yu Kwok, Zhihong Sun, Heping Zhang

**Affiliations:** Key Laboratory of Dairy Biotechnology and Engineering, Ministry of Education, Key Laboratory of Dairy Products Processing, Ministry of Agriculture, Inner Mongolia Agricultural University, Huhhot, P. R. China

**Keywords:** bovine mastitis, PacBio single-molecule real-time sequencing, l6S rRNA, digital droplets PCR, milk microbiota

## Abstract

Mastitis is the most economically important disease of dairy cows. This study used PacBio single-molecule real-time sequencing technology to sequence the full-length of the l6S rRNA from the microbiota in 27 milk samples (18 from mastitis and 9 from healthy cows; the cows were at different stages of lactation). We observed that healthy or late stage milk microbiota had significantly higher microbial diversity and richness. The community composition of the microbiota from different groups also varied greatly. In milk from healthy cows the microbiota was predominantly comprised of *Lactococcus lactis, Acinetobacter johnsonii* and *Bacteroides dorei*, while from mastitis cows it was predominantly comprised of *Bacillus cereus, Clostridium cadaveris* and *Streptococcus suis*. The prevalence of *La. lactis* and *B. cereus* in milk from healthy and mastitis cows was confirmed by digital droplets PCR. Differences in milk microbiota composition could suggest an important role for these microbes in protecting the host from mastitis. Based on the milk microbiota profiles, the Udder Health Index was constructed to predict the risk of bovine mastitis. Application of this predictive model could aid early identification and prevention of mastitis in dairy cows, though the model requires further optimisation using a larger dataset.

## 1 INTRODUCTION

Bovine mastitis is an inflammatory reaction that occurs in the cow’s mammary glands; it is caused either by invasion of pathogenic microbes or by physical/chemical stimulants (Rasmussen, Fogsgaard, Rontved, Klaas, & Herskin, 2011). In general, depending on the degree of inflammation, mastitis can be divided into subclinical and clinical mastitis. Clinical mastitis was diagnosed when the milk from the udder had visible abnormalities (e.g. the presence of floc and/or granules in milk or watery/reduced milk/no milk) or cows showing clinical symptoms, such as elevated body temperature, loss of appetite/refusal to eat, swelling/oedema of the affected quarter (Carolina Espeche et al., 2012; Metzger et al., 2018). In subclinical cases, animals are outwardly healthy, but when the somatic cell count (SCC) of milk samples is greater than 200,000 cells/mL and there is no evidence of clinical infection, they are classified as having subclinical mastitis (Pantoja, Hulland, & Ruegg, 2009). Bovine mastitis is one of the most common and serious diseases in the dairy farming industry; not only does it lead to a decline in milk production and milk quality, but it also increases the elimination and mortality rate of cows. Approximately 33% of the 231 million cows in the world have mastitis, and annual losses due to mastitis are as high as $3.5 billion (Halasa, Huijps, Osteras, & Hogeveen, 2007). Owing to the huge economic significance of bovine mastitis, it is of major global concern.

The etiology of bovine mastitis is complicated and varies amongst countries and farms; it is influenced by environmental/geographical factors, feeding management methods, sanitary conditions, microbial infections, and attributes of individual cows (e.g. age, lactation stage, parity, body type, heredity). In general, the main cause of mastitis is bacterial infection (Haltia, Honkanen-Buzalski, Spiridonova, Olkonen, & Myllys, 2006), as supported by numerous reports of the isolation and identification of mastitis-causing microbes using traditional microbiological methods. However, traditional bacterial culture is relatively slow and laborious and no pathogenic bacteria are detected, using conventional methods, in approximately 25% of milk samples collected from mastitis cows (Taponen, Salmikivi, Simojoki, Koskinen, & Pyorala, 2009). Second-generation sequencing technology only produces short sequence reads with low taxonomic resolution and has failed to identify mastitis-causing microbes (Dohoo et al., 2011; Liao, Lin, & Lin, 2015). Third-generation sequencing technology, such as the PacBio single-molecule real-time (SMRT) sequencing platform, is increasingly being used to characterize the microbiota of environmental samples (e.g. dairy products (Hui et al., 2017), milk (Hou et al., 2015) and food (Nakano et al., 2016) etc. In contrast to the older sequencing technologies, PacBio SMRT sequencing is high throughput and produces long reads capable of microbial identification to species level when used in conjunction with polymerase chain reaction (PCR) of full-length 16S rRNA genes (Mosher et al., 2014).

This study aimed to identify differences in the microbiota of milk from healthy and mastitis cows at different stages of lactation. Milk samples were collected from healthy and mastitis cows from the same dairy. Microbiota profiles in these samples were characterized at the species level by PacBio SMRT sequencing. Abundances of specific microbes were determined using ddPCR. With these data, an Udder Health Index was constructed using the Random Forest Algorithm. Our results serve as a useful reference for designing strategies to prevent and treat mastitis.

## 2 MATERIALS AND METHODS

### 2.1 Experimental design and selection of cows

This present study was conducted at Aoya Modern Ranch in Tai’an City (Shandong Province, China) using 2- to 6-year-old Holstein cows. The average age of calving is 23-76 months, and the milk production is about 3800 kg. From 26 dairy cows with severe clinical mastitis identified by professional veterinarians, 18 were randomly selected as mastitis group. For comparison, 9 milk samples obtained from healthy cows was used, and these cows had no history of mastitis and found to have a SCC lower than 15,000 cells/mL. Milk from the mastitis cows contained obvious floc or granules which prevented measurement of SCC. According to Vijayakumar et al (Vijayakumar et al., 2017). lactation in cows can be divided into early (1 ≤ d ≤ 100), middle (101 ≤ d ≤ 200), and late lactation (201 ≤ d ≤ dry milk period). Dairy cows are managed in accordance with the practices of the herd in the pasture, providing them with green fodder and calculating the amount of concentrated mixture. Furthermore, none of the cows had received antibiotics or any other medication known to influence the microbiota of their milk, in the three months before the study.

### 2.2 Collection of milk samples

Milk samples were collected from 27 cows in total (18 mastitis and 9 healthy cows; 8 early, 5 middle and 14 late lactation stage cows); detailed information was shown in Table S1. Quarter milk samples were collected during the morning milking from all cows in the following way: Udders were thoroughly wiped using a clean dry cloth to remove bedding and visible contaminants; teats were sanitized; two or three streams of foremilk per teat were discarded; a 0.5% iodine solution was applied to the udder; after 60s the iodine was wiped off using another clean dry cloth towel; the collector then put on clean gloves; a further two to three streams of milk per teat were discarded; all teats were scrubbed with 70% isopropanol; a further two streams of milk per teat were discarded after the isopropanol had dried; finally, approximately 10ml were collected into a sterile vial. All raw milk was stored and transported back to the laboratory on ice.

### 2.3 Extraction of metagenomic DNA

DNA was extracted from 3 ml of each raw milk sample using the PowerFood™ Microbial DNA Isolation Kit (MoBio Laboratories, Qiagen, USA) following the manufacturer’s instructions. The quality of the extracted genomic DNA was examined by agarose gel electrophoresis and spectrophotometer analysis (ratio of optical density at 260 nm/280 nm). High quality DNA samples were temporarily stored in the refrigerator at -20°C prior use and for no longer than 6 hours.

### 2.4 Droplet digital PCR

To verify the sequencing results, ddPCR was used to quantify *La. lactis* and *B. cereus*, as they were the most abundant species in milk from healthy and mastitis cows, respectively. Nine samples from each group were included in this analysis. The ddPCR was done using the QX200 system (Bio-Rad, Hercules, CA, USA). Primer 5.0 software was used to design the primers targeting *La. lactis* (lactisF, 5’-AGCAGTAGGGAATCTTCGGCA-3’; lactisR, 5’-GGGTAGTTACCGTCACTTGATGAG-3’) and *B. cereus* (PCERF, 5’-GGATTCATGGAGCGGCAGTA-3’; PCERR3, 5’-GCTTACCTGTCATGGTGTAACTTCA-3’) based on the species-specific genomic region (Francisco Martinez-Blanch, Sanchez, Garay, & Aznar, 2011; Ma et al., 2018). The ddPCR reaction solution was prepared by mixing 2 μl DNA, 0.2 μl each of the upstream and downstream primers, 10 μl 2-fold EvaGreen ddPCRSuperMix and 7.6 μl of sterilized ultra-pure water. The prepared ddPCR reaction solution and the droplet generation oil were added to the droplet generation card and partitioned into 20,000 droplets per sample by the QX200 droplet generator (Cremonesi et al., 2016). The droplets produced by each sample were transferred to a 96-well plate and PCR amplification carried out using the EvaGreen program: 95°C for 10 min, 40-cycles of 94°C for 30 sec, 60°C for *La. lactis* and 62°C for *B. cereus* for 1 min, and 4°C for 5 min, followed by 90°C for 5 min and a hold at 4°C. After thermal cycling, the 96-well plate was loaded into the QX200 droplet reader. Data were collected using QuantaSoft software, and the number of target DNA molecules calculated based on Poisson statistics.

### 2.5 Amplification of full-length 16S rRNA genes and SMRT sequencing

The 16S rRNA genes were amplified by PCR using the purified extracted genomic DNA as templates, with the universal primer pair 27F (5′-AGAGTTTGATCCTGGCTCAG-3′) and 1495R (5′-CTACGGCTACCTTGTTACGA-3′) (Mosher, Bernberg, Shevchenko, Kan, & Kaplan, 2013). At the same time, to identify different samples in the same library, a 16-base identification barcode was added to both ends of all primers. The specific amplification conditions were as follows: 95°C for 3 min, 98°C for 20 s, 60°C for 15 s, 72°C for 30 s, 30 cycles and then 72°C for 2 min.

PCR products from each sample were purified and equal mass-mixed to construct the DNA library for sequencing by the Agilent DNA 1000 Kit, according to the manufacturer’s instructions. The constructed library, sequencing primers and DNA polymerase, were added to the SMRT cells for sequencing using the PacBio RS II instrument.

### 2.6 Bioinformatics processing of high-quality sequences

Quality control of raw data was achieved using the RS_ReadsOfinsert.1 protocol available in the SMRT Portal version 2.3. Specific quality control conditions were based on the following criteria: minimum cycle sequencing number, minimum prediction accuracy, minimum insertion sequence length and maximum insertion sequence length, which were set to 5, 90, 1,400 and 1,800 respectively (Hou et al., 2015). Then all sequences were divided into different samples according to the barcodes on the amplicons. After removal of barcodes and primer sequences, bioinformatics analysis of high-quality sequences was done using the QIIME package (version 1.7), briefly, PyNAST (Caporaso et al., 2010) was used to align the sequences, and UCLUST (Edgar, 2010) was done under 100% clustering of sequence identity to obtain representative sequences. Afterwards, sequences were classified into operational taxonomic units (OTU) according to 97% similarity. Chimeric OTU sequences were detected and removed using ChimeraSlayer (Haas et al., 2011). The Ribosomal Database Project (RDP, Release 11.5) (Cole et al., 2007), Greengenes (version 13.8) (DeSantis et al., 2006) and Silva (version 128) (Quast et al., 2013) databases were used to assign the taxonomy of each OTU representative sequence with an 80% confidence threshold (Hou et al., 2015). A *de novo* taxonomic tree was constructed for representative OTU sets using FastTree for downstream analysis (Price, Dehal, & Arkin, 2009). The alpha diversity of each sample was assessed based on the sample with the lowest sequencing depth. All sequencing data generated have been uploaded to the MG-RAST database under the project number mgp88495 (http://www.mg-rast.org).

### 2.7 Random Forest algorithm for predicting cow udder health

Here a mastitis prediction model was built to predict the udder health status of dairy cows based on the default settings of the Random Forest machine learning algorithm available in R package (3.1.1) (Cutler, Edwards, Beard, Cutler, & Hess, 2007). Briefly, the prediction was based on regression of the milk microbiota profiles of all 27 cows against their mastitis health status. The Random Forest algorithm ranked all species into ‘character importance’ and used the ‘rfcv’ function to generate more than 100 predictions to determine the number of top discriminant species required for calculating the Udder Health Index. Then the Udder Health Index was calculated based on the relative abundance of the predicted top discriminatory milk microbes.

### 2.8 Statistical analyses

The R software package was used for statistical analysis (http://www.rproject.org/). Wilcoxon and Kruskal-Wallis tests were used to evaluate differences in microbial between groups with a cut-off confidence level of 95%. Principal coordinate analysis (PCoA) was used to observe the distribution of samples. Permutational multivariate analysis of variance (PERMANOVA) (Anderson & Walsh, 2013) was done on the different groups of samples in the R language ‘vegan’ (Nilsson et al., 2010) package to reveal the effect of different groups on the target microbiota of healthy and mastitis cows. The ‘ggplot2’ package (Ginestet, 2011) was used to analyze and visualize the raw results obtained by QIIME. Cytoscape 3.5.1 was used for network building.

## 3 RESULTS

### 3.1 Sequence coverage and alpha diversity of microbiota from milk

A total of 169,308 high-quality 16S rRNA sequences were produced from the 27 samples included in this study (average = 6,270; range = 1,010-15,174; SD = 3,905). After PyNAST alignment and UCLUST classification at 100% similarity, 82,592 representative sequences remained. Upon removal of chimeric sequences, they were classed into 6,850 OTUs for downstream analysis.

All Shannon diversity curves leveled off, suggesting that the sequencing depth was sufficient to capture representative microbial populations present in the samples (Figure S1), although new phylotypes may still be found with further sequencing. The microbial abundance and species richness in each sample were assessed by the number of observed OTUs and the Shannon diversity index, respectively. Both the number of observed OTUs and Shannon diversity index were significantly higher in healthy compared with mastitis cows (Figure 1a, b; *P* < 0.001). We also found that both indices were significantly higher during late compared with early lactation, while no significant difference was observed amongst other lactation stages (Figure 1c, d; *P* < 0.05).

**Figure 1.**
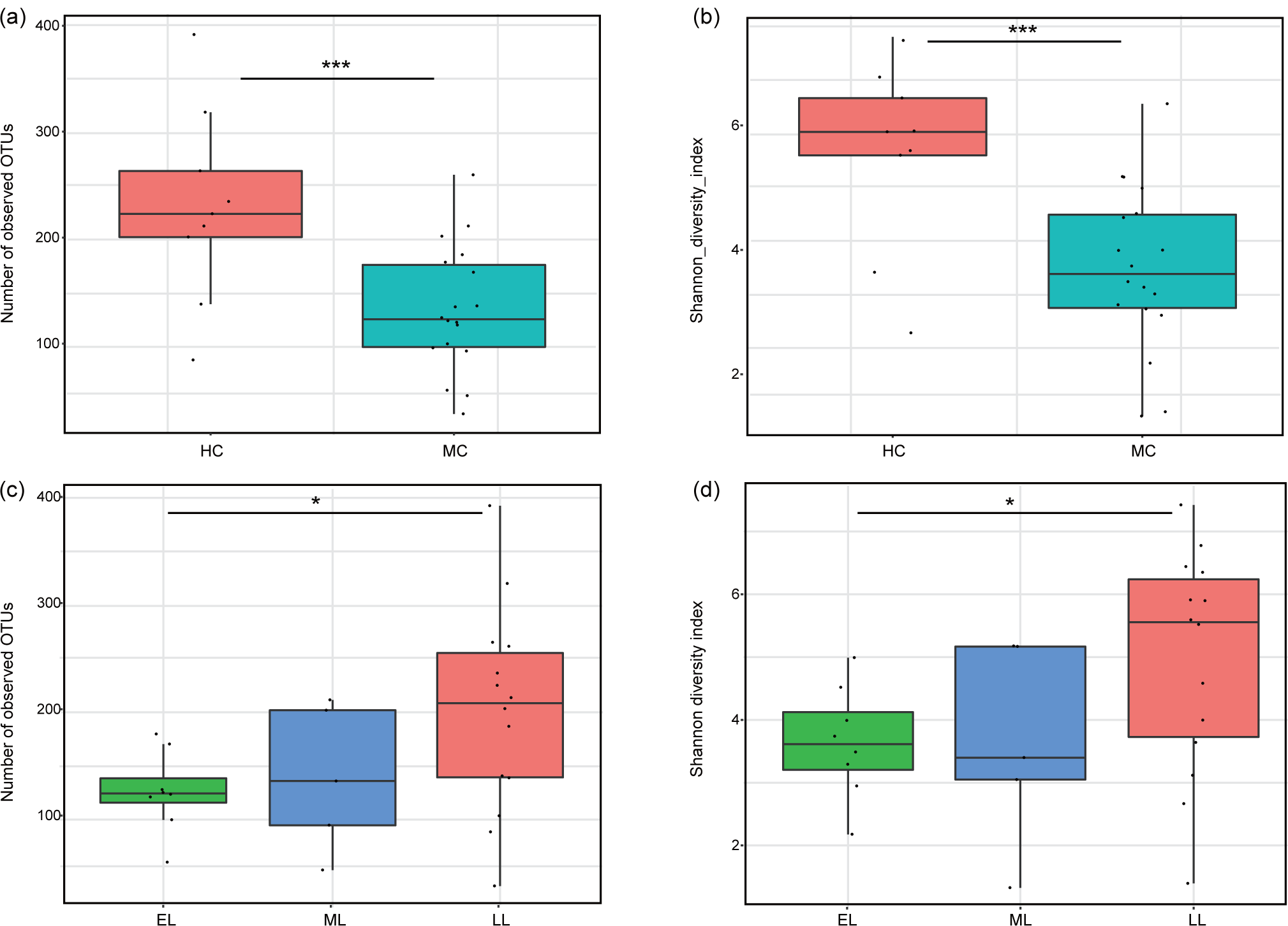
Comparison of the number of observed OTUs and Shannon diversity index for the microbiota in milk. The healthy and mastitis cows (a, b); early, middle and late lactation (c, d). Healthy and mastitis cows are represented by ‘HC’ and ‘MC’, respectively. Early, middle, and late lactation stages are represented by ‘EL’, ‘ML’ and ‘LL’, respectively. Single and triple asterisks represent *P* < 0.05 and *P* < 0.001, respectively.

### 3.2 Identity of microbiota in milk

*Firmicutes* (73.0%) and *Bacteroidetes* (10.9%) were the two main bacterial phyla. At the genus level, there were 12 genera with an average relative abundance of more than 1% (Figure 2a), including *Bacillus* (28.5%), *Clostridium* (10.6%), *Lactobacillus* (10.4%), *Lactococcus* (7.9%) and *Bacteroides* (6.1%). At the species level, 15 of which had an average relative abundance of over 1% (Figure 2c), including *B. cereus* (27.6%), *La. lactis* (7.8%), *Lactobacillus helveticus* (6.3%), *Clostridium limosum* (4.4%), *Helcococcus ovis* (3.4%), and *Clostridium cadaveris* (3.2%) et al.

**Figure 2.**
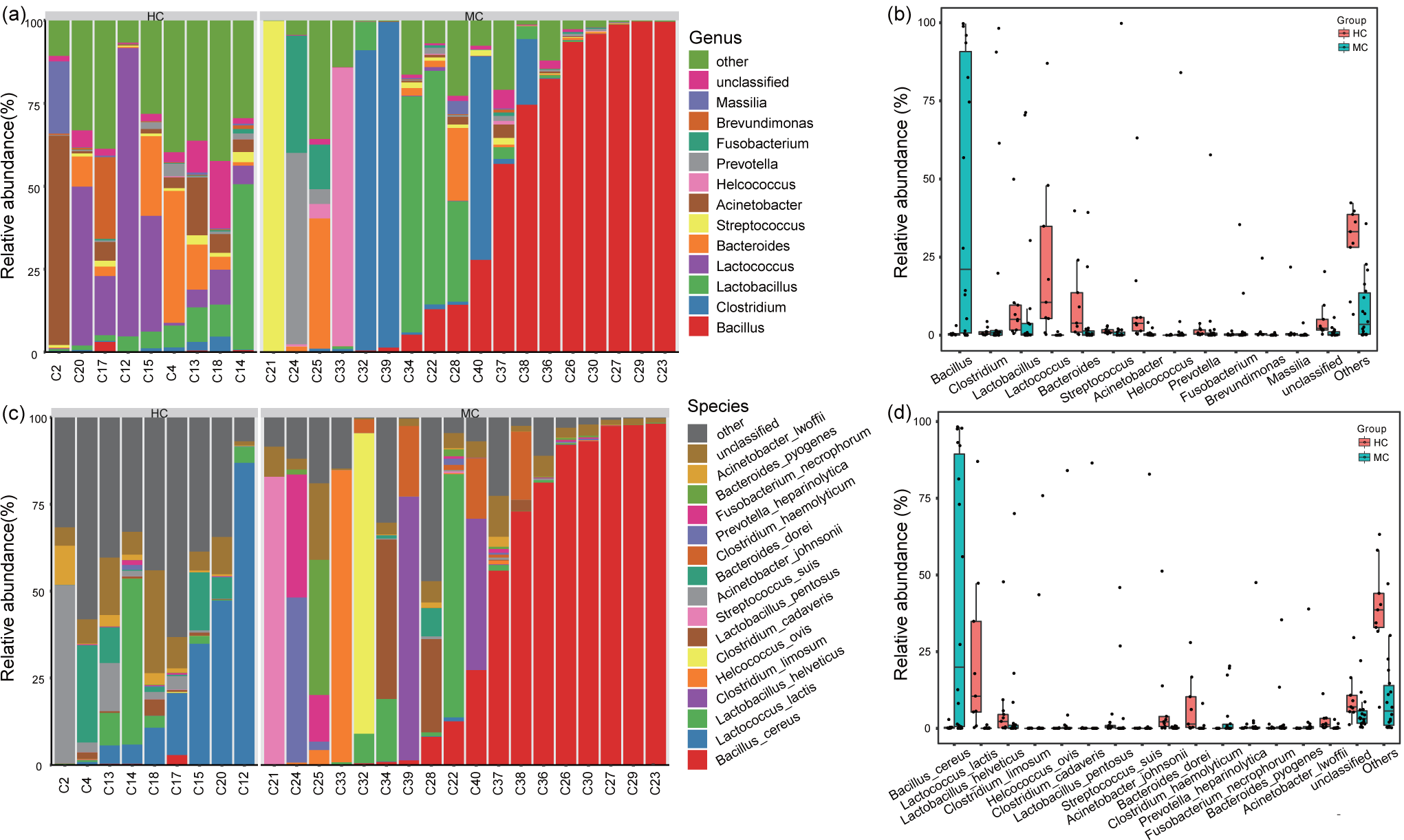
Community composition of the bacterial microbiota in milk from healthy and mastitis cows. Milk bacterial community at the genus (a and b) and species (c and d) levels. Healthy and mastitis cows are represented by ‘HC’ and ‘MC’, respectively.

The mean microbiota composition varied largely between groups. Whereas the horizontal lines in the box-plots indicate the median, we found that more *Lactobacillus, Lactococcus, Bacteroides* and *Acinetobacter* were detected in healthy cows, while more *Bacillus, Clostridium* and *Streptococcus* were observed in mastitis cows (Figure 2b). Significantly more *La. lactis* was found in the milk of healthy cows, while *B. cereus* was found in the mastitis cows (*P* < 0.05, Figure 2d). The relative abundance of *Bacillus* and *Clostridium* were not significantly different in milk collected during the early and middle compared with late lactation. In addition, significantly more *La. lactis, S. suis, A. johnsonii*, and *B. dorei* were detected in milk from late compared with early and middle lactation cows (*P* < 0.05). The relative abundance of *H. ovis* and *P. heparinolytica* gradually decreased as lactation day increased (Figure S2).

### 3.3 Comparative analysis of community structure in milk from cows

A PCoA was done based on the Bray Curtis distance to visualize differences in the milk microbiota between the healthy and mastitis groups. Two distinct clusters formed on the PCoA score plot, representing milk samples collected from healthy and mastitis cows, respectively (Figure 3a). This suggests large difference in the microbiota community structure and compositon between the two groups. Results of the PERMANOVA test revealed that mastitis was a significant factor contributing to differences in microbiota between the two groups (*P* = 0.001).

**Figure 3.**
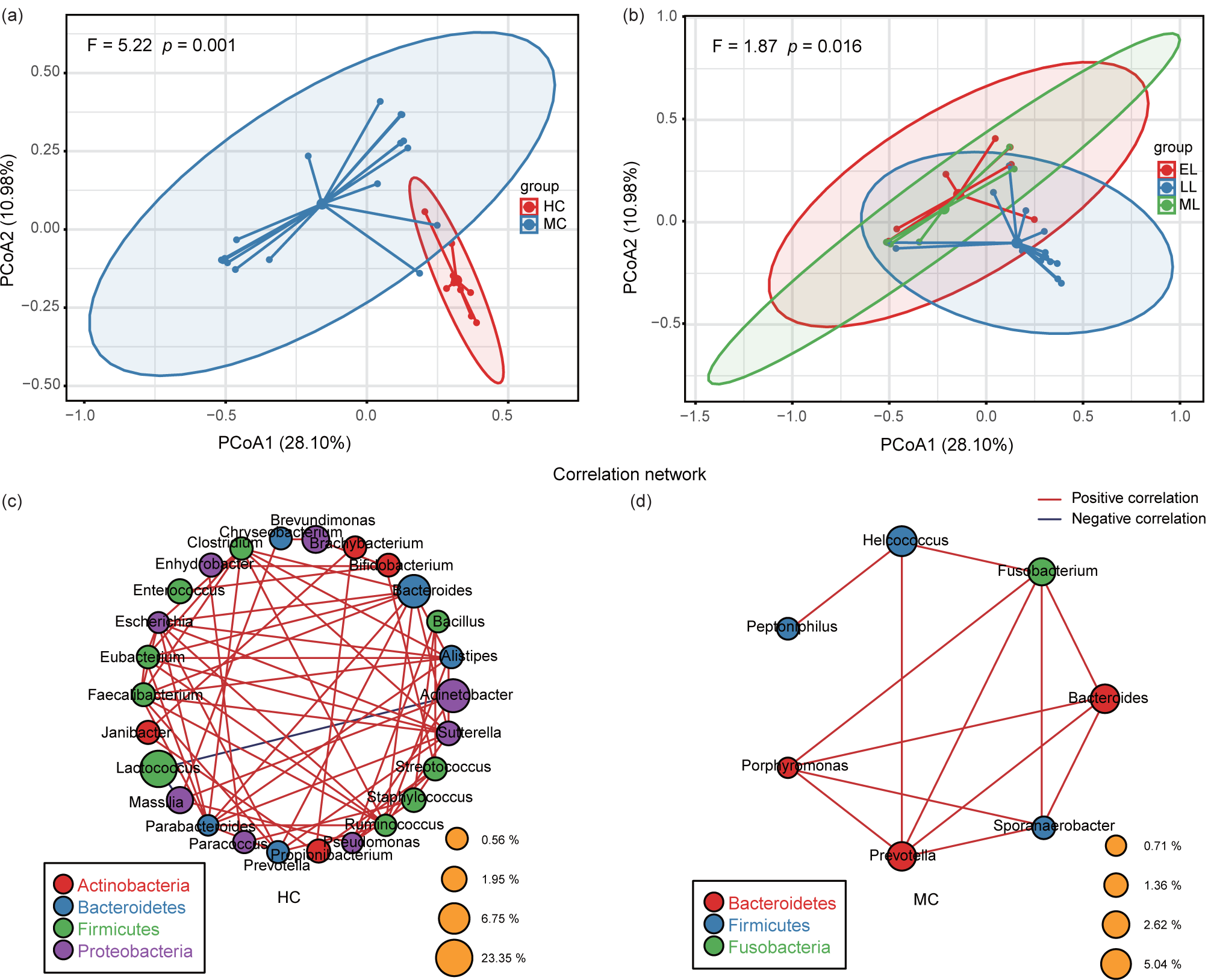
Principal coordinates analysis (PCoA) and correlation networks of the bacterial microbiota in milk from healthy and mastitis cows. They are grouped based on mastitis status (a) and lactation stage (b). The co-occurrence relationship was calculated using Spearman’s rank correlation analysis of the microbiota in milk from healthy (c) and mastitis (d) cows. Only major genera of average relative abundance > 0.5% are included in the analysis. The diameter of the circles represents the abundance of the genera. Healthy and mastitis cows are represented by ‘HC’ and ‘MC’, respectively. Early, middle, and late lactation stages are represented by ‘EL’, ‘ML’ and ‘LL’, respectively.

When samples were grouped based on the three lactation stages, the distinction between groups were less clear. Clusters associated with early and middle lactation milk overlapped each other and were separate from the cluster representing late lactation milk. This indicates that the former two groups were more similar to each other than they were to the latter group (Figure 3b). The PERMANOVA test revealed significant difference in the milk microbiota composition of the three lactation stages (*P* = 0.016). However, the effect size of lactation stage on milk microbiota was smaller than that of mastitis status. It is worth noting that most late lactation dairy cows were healthy.

Wilcoxon and Kruskal-Wallis tests were done to identify differentially abundant taxa in the different groups. Species that were significantly differentially abundant (average relative abundance > 0.5%) are listed in Table S2 and S3. Significantly more *La. lactis, A. johnsonii* and *B. dorei* were found in the milk of healthy (*P* < 0.01) compared with mastitis cows, while significantly more *B. cereus* and *S. suis* were found in the milk of mastitis compared with healthy cows (*P* < 0.05). Meanwhile, significantly more *La. lactis, A. johnsonii, B. dorei, S. stercoricanis* and *S. suis* were present in the milk of late lactation cows compared with other lactation stages (*P* < 0.05); the two latter species were found only during late lactation.

### 3.4 Correlation analysis of microbiota community structure in milk from cows

Spearman correlation analyses were done to identify co-occurrence relationships amongst the major bacterial genera (an average relative abundance > 0.5%) in milk from healthy and mastitis cows. The results are expressed as correlation network diagrams. The milk microbiota of the healthy group seemed to be closely interrelated, contrasting with the overall weak networking apparent amongst bacterial genera in milk from mastitis cows. In milk from healthy cows, *Lactococcus* was significantly and negatively correlated with *Acinetobacter* (r = -0.78; *p* = 0.01) and *Massilia* (r = -0.70; *p* = 0.04, Figure 3c), respectively. It is interesting to note that the *Faecalibacterium* formed the highest number of significant correlations with other genera from milk. In contrast, *Faecalibacterium* didn’t correlate with any genera in the milk from mastitis cows (Figure 3d).

### 3.5 Quantification of microbial in cows milk by ddPCR and construction of Udder Health Index

To verify the sequencing results, ddPCR was done to quantify the number of *La. lactis* and *B. cereus*, as they were the most abundant species in the healthy and mastitis groups, respectively. For the healthy group, there were significantly more *La. lactis* (2.16×103 copies/ml) compared with *B. cereus* (*P* < 0.01), and *B. cereus* was only detected in the sample C17 (healthy group). An opposite trend was observed for the mastitis group, with a significantly higher abundance of *B. cereus* (3.25×104 copies/ml) compared with *La. lactis* (9.00×102 copies/mL; *P* < 0.001) (Figure 4a). These results were consistent with those found by DNA sequencing.

**Figure 4.**
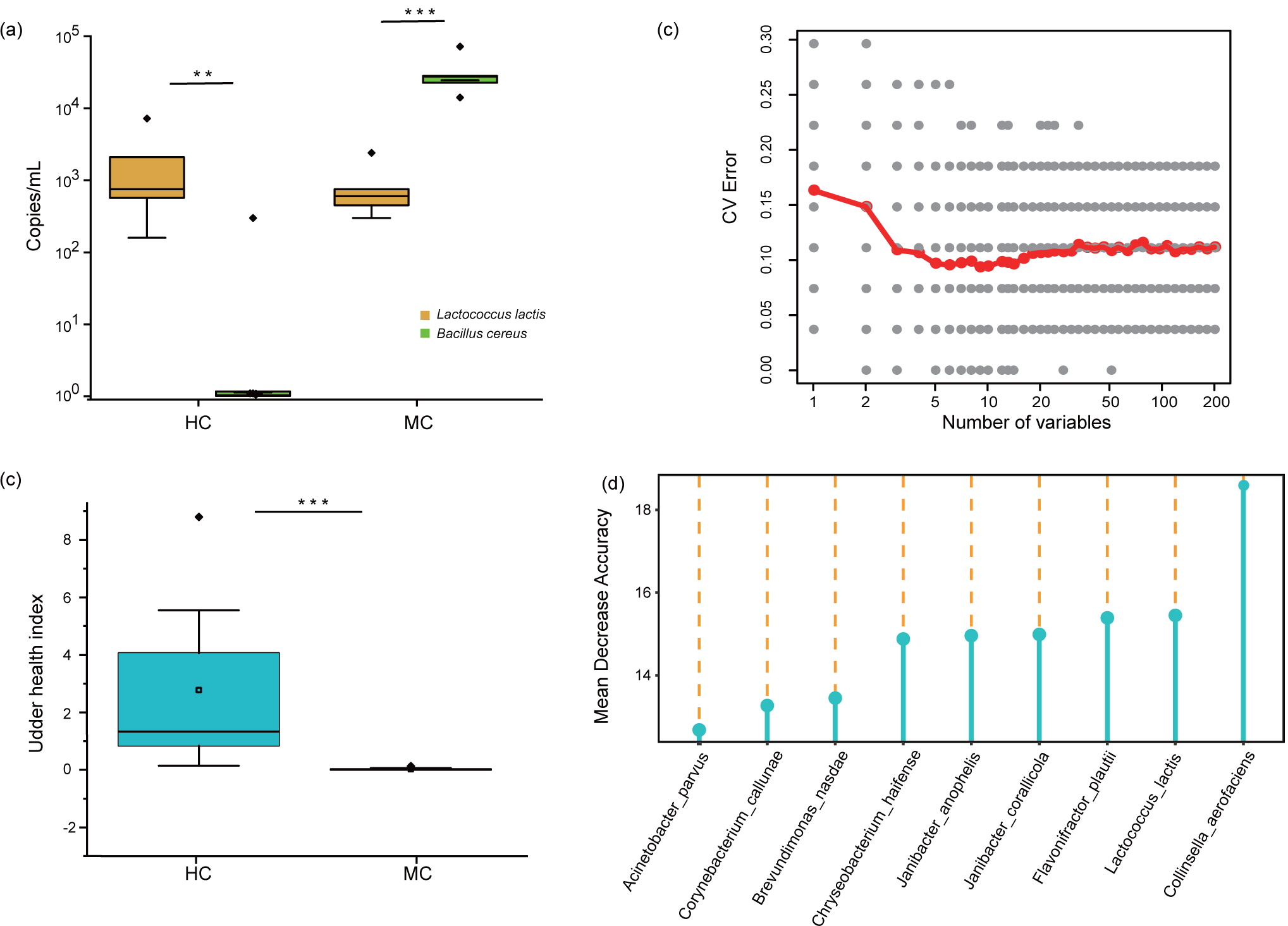
Quantification of *La. lactis* and *B. cereus* in cows’ milk by ddPCR and construction of Udder Health Index. (a) The prevalence of *La. lactis* and *B. cereus* in milk from healthy and mastitis cows. (b) Model marker species selected based on minimum CV error. (c) The nine selected marker species of the model. (d) Udder Health Indices in the healthy and mastitis groups. Healthy and mastitis cows are represented by ‘HC’ and ‘MC’, respectively. Single and triple asterisks represent *P* < 0.05 and *P* < 0.001, respectively.

The Random Forest regression model was applied to predict the udder health of cows. The relative abundances of bacterial species detected in milk were regressed against each cow’s health status. The top nine mastitis-discriminatory marker species were selected based on the minimum ‘CV error’, and an Udder Health Index was constructed (Figure 4b,d). Udder Health Indices for the healthy and mastitis groups were calculated; a higher value represented a healthier status. As expected, the Udder Health Index was significantly higher for the healthy compared with the mastitis group (*P* < 0.001, Figure 4c). To verify the accuracy of the constructed model, the SCC of the healthy milk samples was measured (Table 1). But this was not possible for the mastitis group due to the milk flocs were obvious and the granules were large, accompanied by blood and different degree of coagulation. However, the SCC associated well with the predicted Udder Health Index of the healthy cows, validating the current model.

**Table 1.**
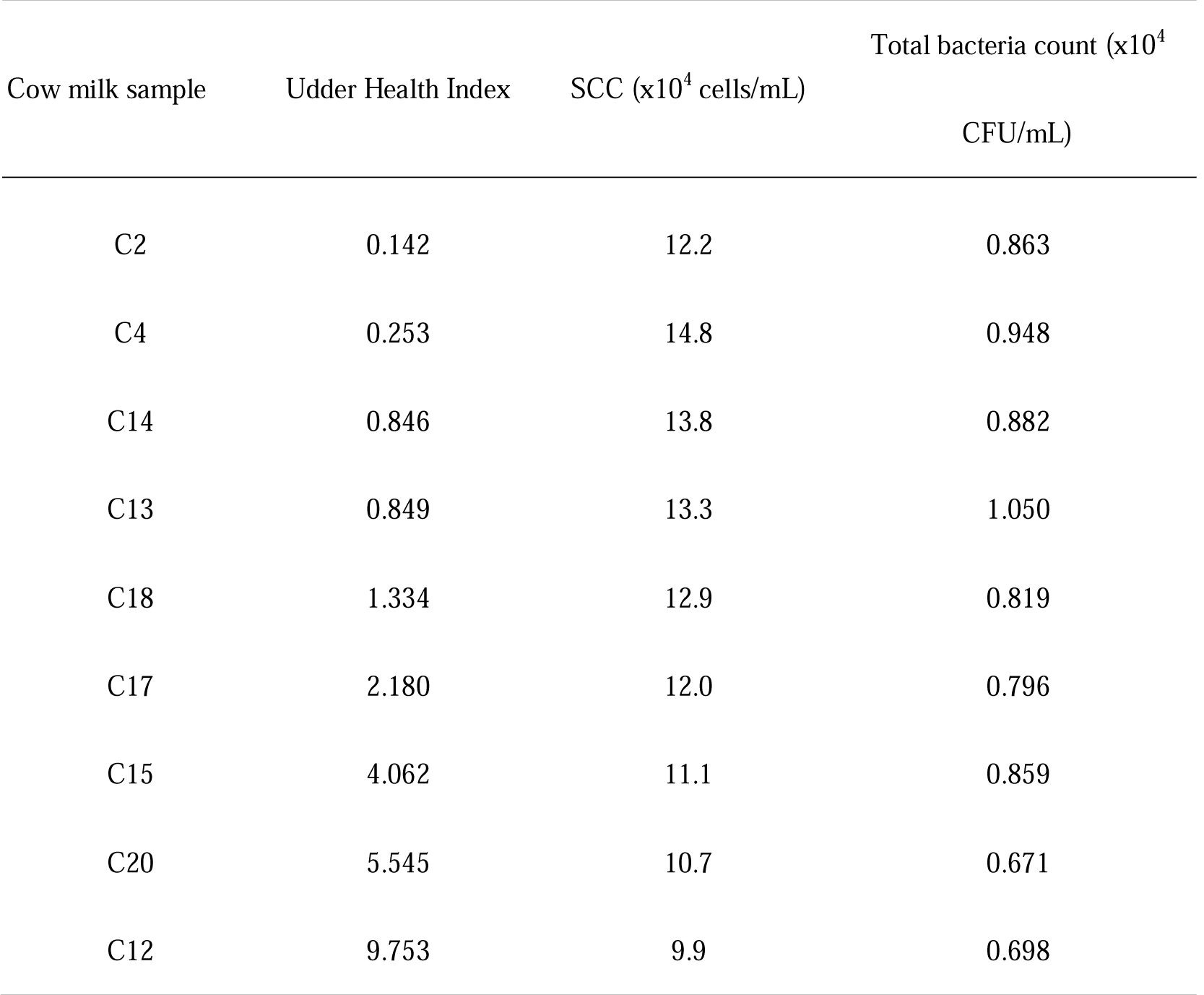
Udder Health Index, somatic cell count (SCC), and total bacterial count of cow milk samples

## 4 DISCUSSION

Mastitis is a common disease among dairy cows; symptoms include shortening of lactation period and milk production, increase in leukocytes in the milk, and lesions in mammary tissues (Paster, Dewhirst, Olsen, & Fraser, 1994). Since bovine mastitis results in huge economical losses, its prevention and treatment have attracted wide attention. Bovine mastitis can also increase the microbial load in milk resulting in rapid changes in the quality and shelf life of raw milk and related products (Murphy, Martin, Barbano, & Wiedmann, 2016). Pathogenic bacteria in raw milk from mastitis cows are a serious food safety issue as they may lead to human disease. In addition, clinical mastitis is a serious animal welfare concern because mastitis is debilitating and painful (Fromm & Boor, 2004; Halasa et al., 2007). Milk is produced by the mammary tissues of lactating cows; the appearance, texture, and quantity of secreted cow milk are indicative of the health of the mammary tissue and thus serve as indicators for clinical mastitis. This study used PacBio SMRT sequencing technology to reveal the community composition of microbiota in milk from dairy cows, and identified differences in the milk microbiota of healthy and mastitis cows, and cows at different stages of lactation. We observed a significantly higher microbial diversity and richness in milk from healthy compared with mastitis cows, as reported in other studies (Braem et al., 2012; Kuehn et al., 2013). Our data also showed that the microbial diversity and richness of milk also increased significantly during late lactation. On the other hand, the increase in microbial diversity could indicate a healthy state of the cows, as the mastitis milk might be dominated by certain pathogens, which would be reflected by the diminished microbiota diversity. Moreover, changes in milk microbiota at different lactation stages could be associated with changes in the nutritional content of the milk, e.g. the total concentration of milk oligosaccharides was found to decrease in the early and mid lactation stages. The anionic oligosaccharides including N-glycolylneuraminic acid decreased more rapidly than the neutral oligosaccharides in lactation (Nakamura et al., 2003; Tao, DePeters, German, Grimm, & Lebrilla, 2009). Therefore, when cows suffer from mastitis, the mammary tissues may be overwhelmed with harmful bacteria; this could induce localized immune responses that suppress the healthy resident microbiota and reduce microbial diversity and richness in the mammary gland (Kuang et al., 2009).

We found that the distribution of bacterial taxa present in milk also varied greatly between healthy and mastitis cows, as found in other studies (Falentin et al., 2016; Oikonomou, Machado, Santisteban, Schukken, & Bicalho, 2012). The predominant genera detected in this work included *Lactobacillus, Streptococcus, Acinetobacter* and *Bacillus*, which are known to be common in milk (N. Li et al., 2018). We found significantly more *Lactobacillus, Lactococcus* and *Acinetobacter* in milk from healthy compared with mastitis cows. The relative abundance of *Lactobacillus* in milk is known to be negatively associated with milk SCC, an indicator of the seriousness of mastitis; thus, these bacteria are crucial in maintaining the health of the mammary tissues and suppressing local infection and inflammation (Yu, Ren, Xi, Huang, & Zhang, 2017). In contrast, *Lactococcus* is known to be able to cause bovine mastitis (Rodrigues, Lima, Higgins, Canniatti-Brazaca, & Bicalho, 2016). *Acinetobacter* is widely distributed in nature and is often detected in milk, soil and water. It is also considered as an opportunistic pathogen associated with wounds and skin infections (Dortet, Legrand, Soussy, & Cattoir, 2006; L. Li et al., 2016). Results of the Spearman correlation analysis in our study indicated a negative correlation between *Lactococcus* and *Acinetobacter*, suggesting that there may be a competition inhibition relationship between them. *Bacteroides* are typical milk bacteria that may also have a role in maintaining healthy mammary tissues (Quigley et al., 2013). Differences in the correlation patterns between the milk microbiota from healthy and mastitis cows may suggest an important role of milk microbiota in protecting cows from mastitis. Our data revealed an increase in the relative abundances of *Bacillus, Clostridium* and *Streptococcus* in milk from mastitis compared with healthy cows. *Staphylococcus* has been reported as the most common cause of mastitis in cows worldwide, followed by *Streptococcus* (Moroni et al., 2006). *Clostridium* species have been reported to cause abscesses in mammary tissues of sows and humans as well as gangrene in cows (Durojaiye, Gaur, & Alsaffar, 2011; Osman, El-Enbaawy, Ezzeldeen, & Hussein, 2009). The *Bacillus* genus includes some important causative agents of mastitis, e.g. *B. cereus* (Parkinson, Merrall, & Fenwick, 1999); in this study *B. cereus* increased in relative abundance (up to 27.55%) in milk from mastitis compared with healthy cows. By ddPCR, we confirmed the elevated abundance of *Bacillus cereus* in the mastitis group (3.25×104 copies/mL); it was practically absent in the healthy group. These results suggest a possible link between *B. cereus* and bovine mastitis.

Another spectrum of mastitis-associated bacterial sequences detected in this study was the anaerobes, including *F. necrophorum* and *B. dorei*. Although these bacteria are unlikely to be causative agents of clinical bovine mastitis, they are known to be associated with summer mastitis and may interact with other pathogens such as *Trueperella pyogenes* (Oikonomou et al., 2012; Pyorala, Jousimies-Somer, & Mero, 1992). Significantly more *P. heparinolytica* and *H. ovis* sequences were detected in milk from cows at the early lactation stage in this study, both *P. heparinolytica* and *H. ovis* are potential pathogens that have previously been isolated from mammary gland wounds causing localized infection (Paster et al., 1994). It is suspected that the presence of *H. ovis* in cows milk indicates involvement in the pathogenesis of mastitis (Schwaiger et al., 2012).The relative abundances of *La. lactis, A. johnsonii* and *B. dorei* were higher during late lactation compared with the early and middle lactation stages. *B*. *dorei* can suppress the production of lipopolysaccharides by intestinal gut microbes and thus reduces pro-inflammatory immune responses (Yoshida et al., 2018). Furthermore, changes in susceptibility to mastitis might be coupled with changes in community composition of the milk microbiota, such as increases in *B. dorei* and *La. lactis* (Xu et al., 2017).

Cows suffering from subclinical mastitis can easily develop clinical mastitis if they are not spotted early enough and managed appropriately. Therefore, it is important to predict the likely risk of mastitis developing. Current practice defines subclinical mastitis using a cut-off SCC threshold level of 200 000 cells/mL. However, the determination of milk SCC is widely used to monitor udder health, but there are still some limitations: (1) it may take a longer time for the SCC to return to that of a healthy state after pathogen clearance, so there is a window period when the diagnosis based on SCC level would not be accurate; (2) the SCC value fluctuates largely with individual milk yield and other environmental factors; (3) diagnosis based purely on SCC is mainly applicable to mastitis cows caused by contagious pathogens, while mastitis caused by environmental factors would be hard to detect due to the relatively small changes in SCC (Pyorala et al., 1992; Sharma, Singh, & Bhadwal, 2011). In this study we used the Random Forest model identified the 9 top marker species that were indicative of bovine mastitis, and constructed an Udder Health Index based on the relative abundances of these species. We validated the model by associating the index with the SCC of healthy cows. Our results showed that the accuracy of the model was 88.89%. Thus, this index would be a good complementary indicator that helps detect early changes in cow health. One limitation of the current model is the small sample size, therefore, larger scale future works will be necessary to optimize and verify this model.

## 5 CONCLUSION

The results of this study showed great variation in the microbiota in milk from healthy and mastitis cows. Overall there was a high relative abundance of commensals in healthy milk and a high relative abundance of potential pathogens in milk from mastitis cows. Finally, the Udder Health Index constructed is helpful for early identification of mastitis risk and has the potential to be used to prevent mastitis from developing, and prompt measures should be taken to prevent further development of clinical mastitis.

## ACKNOWLEDGEMENTS

This research was supported by the China Agriculture Research System (Grant CARS-36) of the Inner Mongolia Science & Technology Projects. The authors wish to sincerely thank Mr. Yang Ku for allowing the collection of milk samples from the cows on his farm. We also thank Dr. Lai Yu-Kwok for their valuable revision of the English text.

