## Supplementary material for "Microbiota in milk from healthy and mastitis cows varies greatly in diversity, species richness and composition, as revealed by PacBio sequencing": Supporting information.docx

**Table S1** cow information

| Cow id | Age (days) | Number of parity | | Lactation days | Group | Lactation stage |
| --- | --- | --- | --- | --- | --- | --- |
| C2 | 2303 | | 4 | 308 | Healthy | Late |
| C4 | 988 | | 1 | 271 | Healthy | Late |
| C12 | 1122 | | 1 | 437 | Healthy | Late |
| C13 | 1067 | | 1 | 382 | Healthy | Late |
| C14 | 938 | | 1 | 260 | Healthy | Late |
| C15 | 2201 | | 4 | 256 | Healthy | Late |
| C17 | 1152 | | 1 | 318 | Healthy | Late |
| C18 | 1309 | | 2 | 275 | Healthy | Late |
| C20 | 2304 | | 4 | 355 | Healthy | Late |
| C21 | 2343 | | 6 | 232 | Mastitis | Late |
| C22 | 2347 | | 4 | 311 | Mastitis | Late |
| C23 | 2269 | | 5 | 43 | Mastitis | Early |
| C24 | 2219 | | 5 | 83 | Mastitis | Early |
| C25 | 1956 | | 4 | 188 | Mastitis | Middle |
| C26 | 1877 | | 4 | 144 | Mastitis | Middle |
| C27 | 1739 | | 3 | 250 | Mastitis | Late |
| C28 | 1645 | | 2 | 363 | Mastitis | Late |
| C29 | 1581 | | 3 | 200 | Mastitis | Middle |
| C30 | 1456 | | 3 | 40 | Mastitis | Early |
| C32 | 1197 | | 2 | 103 | Mastitis | Middle |
| C33 | 1147 | | 2 | 55 | Mastitis | Early |
| C34 | 1139 | | 2 | 39 | Mastitis | Early |
| C36 | 1061 | | 1 | 320 | Mastitis | Late |
| C37 | 775 | | 1 | 101 | Mastitis | Middle |
| C38 | 747 | | 1 | 69 | Mastitis | Early |
| C39 | 692 | | 1 | 5 | Mastitis | Early |
| C40 | 665 | | 1 | 3 | Mastitis | Early |

**Table S2** Milk bacterial microbiota composition of healthy and mastitis

| Species | Relative abundance (%) | | Median, range (%) | | P-value |
| --- | --- | --- | --- | --- | --- |
|  | Healthy | Mastitis | Healthy | Mastitis |  |
| *Lactococcus lactis* | 23.265 | 0.095 | 2.127, 0.063-51.214 | 0.0123, 0-0.804 | 1.58E-03 |
| *Acinetobacter johnsonii* | 8.475 | 0.162 | 1.212, 0.195-11.286 | 0.021, 0-2.888 | 0.005 |
| *Bacteroides dorei* | 7.035 | 0.501 | 0.123, 0-2.871 | 19.942, 0-98.246 | 0.002 |
| *Brevundimonas diminuta* | 2.494 | 0.045 | 1.387, 0.059-27.952 | 0, 0-8.117 | 0.036 |
| *Acinetobacter lwoffii* | 2.459 | 0.295 | 0.14, 0-2.552 | 0, 0-2.057 | 0.005 |
| *Massilia aurea* | 2.395 | 0.008 | 0.533, 0.074-1.778 | 0, 0-0.311 | 0.003 |
| *Sutterella stercoricanis* | 1.552 | 0.025 | 0.033, 0-22.201 | 0, 0-0.642 | 0.001 |
| *Gossypium hirsutum* | 1.339 | 0.008 | 0, 0-0.223 | 0, 0-86.459 | 0.049 |
| *Enterococcus durans* | 1.026 | 0.179 | 0.64, 0.348-1.192 | 0, 0-1.925 | 0.005 |
| *Streptococcus thermophilus* | 0.996 | 0.182 | 0.46, 0.059-5.979 | 0, 0-2.1 | 0.007 |
| *Escherichia coli* | 0.877 | 0.087 | 0.209, 0.01-3.383 | 0, 0-1.29 | 0.005 |
| *Ruminococcus bromii* | 0.859 | 0.034 | 0.19, 0-3.352 | 0, 0-0.561 | 0.002 |
| *Staphylococcus epidermidis* | 0.774 | 0.076 | 0.051, 0-11.472 | 0, 0-0.111 | 0.003 |
| *Enhydrobacter aerosaccus* | 0.736 | 0.182 | 0.578, 0.123-1.58 | 0, 0-0.198 | 0.005 |
| *Faecalibacterium prausnitzii* | 0.717 | 0.034 | 0.539, 0-4.634 | 0, 0-45.886 | 0.005 |
| *Propionibacterium acnes* | 0.699 | 0.104 | 10.503, 0.237-86.974 | 0.004, 0-1.124 | 0.003 |
| *Pseudomonas stutzeri* | 0.694 | 0.008 | 0.063, 0-20.72 | 0, 0-0.107 | 0.036 |
| *Prevotella copri* | 0.679 | 0.030 | 0.38, 0-2.706 | 0, 0-0.954 | 0.012 |
| *Bacteroides uniformis* | 0.658 | 0.128 | 0.094, 0-3.536 | 0, 0-0.281 | 0.005 |
| *Staphylococcus chromogenes* | 0.635 | 0.027 | 0.584, 0.296-1.569 | 0.011, 0-0.58 | 0.008 |
| *Janibacter anophelis* | 0.632 | 0.030 | 0.06, 0-5.646 | 0, 0-0.104 | 0.001 |
| *Parabacteroides merdae* | 0.630 | 0.065 | 0.335, 0.02-3.229 | 0, 0-0.524 | 0.016 |
| *Brachybacterium faecium* | 0.550 | 0.041 | 0.053, 0-2.679 | 0, 0-0.422 | 0.003 |
| *Bacillus cereus* | 0.446 | 41.102 | 0.578, 0.105-1.931 | 0, 0-0.525 | 0.044 |
| *Streptococcus suis* | 0.034 | 4.628 | 0.584, 0.256-2.437 | 0.052, 0-1.216 | 0.019 |
| *Clostridium haemolyticum* | 0.000 | 3.534 | 0.627, 0-6.888 | 0, 0-0.449 | 0.032 |

**Table S3** Milk bacterial microbiota composition of different lactation stages

| Species | Relative abundance (%) | | | Median, range (%) | | | P-value |
| --- | --- | --- | --- | --- | --- | --- | --- |
|  | Early | Middle | Late | Early | Middle | Late |  |
| *Lactobacillus paracasei* | 0.58 | 0 | 0.27 | 0, 0-0.311 | 0.017, 0-0.642 | 0.713, 0-51.214 | 0.031 |
| *Acinetobacter johnsonii* | 0.05 | 0.16 | 5.57 | 0.005, 0-0.193 | 0.019, 0-2.888 | 0.534, 0-11.286 | 0.034 |
| *Acinetobacter lwoffii* | 0.03 | 0.61 | 1.72 | 0, 0-0.898 | 0, 0-0 | 0.339, 0-27.952 | 0.003 |
| *Bacteroides dorei* | 0.11 | 0 | 5.1 | 0, 0-0.014 | 0, 0-0 | 0.093, 0-7.687 | 0.003 |
| *Bacteroides plebeius* | 0 | 0 | 0.62 | 0, 0-0.249 | 0, 0-0 | 0.102, 0-2.552 | 0.043 |
| *Bacteroides uniformis* | 0.03 | 0 | 0.57 | 0, 0-0 | 0, 0-0.642 | 0.032, 0-22.201 | 0.032 |
| *Brevundimonas diminuta* | 0 | 0.13 | 1.61 | 0.004, 0-1.290 | 0, 0-0 | 0.062, 0-3.383 | 0.039 |
| *Escherichia coli* | 0.18 | 0 | 0.57 | 0, 0-0.048 | 0, 0-0 | 0.13, 0-3.352 | 0.045 |
| *Faecalibacterium prausnitzii* | 0.01 | 0 | 0.5 | 0, 0-4.415 | 0, 0-0 | 0.081, 0-0.861 | 0.034 |
| *Lactococcus lactis* | 0.03 | 0.03 | 15.05 | 0.009, 0-0.173 | 0, 0-0.116 | 3.225, 0-86.974 | 0.008 |
| *Propionibacterium acnes* | 0.03 | 0.13 | 0.52 | 0.011, 0-0.159 | 0, 0-0.535 | 0.511, 0-1.569 | 0.008 |
| *Ruminococcus bromii* | 0.01 | 0 | 0.59 | 0, 0-0.044 | 0, 0-0 | 0.107, 0-3.229 | 0.027 |
| *Staphylococcus epidermidis* | 0.07 | 0.09 | 0.53 | 0, 0-0.525 | 0, 0-0.428 | 0.352, 0-1.931 | 0.031 |
| *Streptococcus suit* | 0 | 0 | 5.97 | 0, 0-0 | 0, 0-0 | 0.015, 0-82.816 | 0.007 |
| *Streptococcus thermophilus* | 0.17 | 0.15 | 0.72 | 0.022, 0-1.216 | 0.052, 0-0.642 | 0.513, 0-2.437 | 0.043 |
| *Sutterella stercoricanis* | 0 | 0 | 1.03 | 0, 0-0 | 0, 0-0 | 0.078, 0-6.888 | 0.03 |

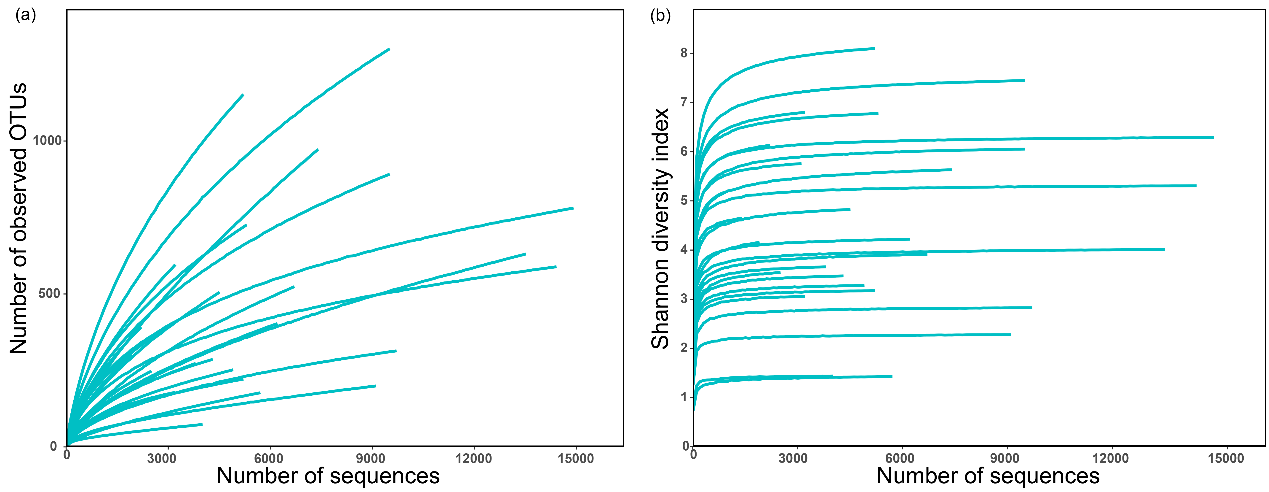

**Figure S1.** Alpha-diversity of dairy milk microflora. (a) Number of observed OTUs; (b) Shannon diversity index.

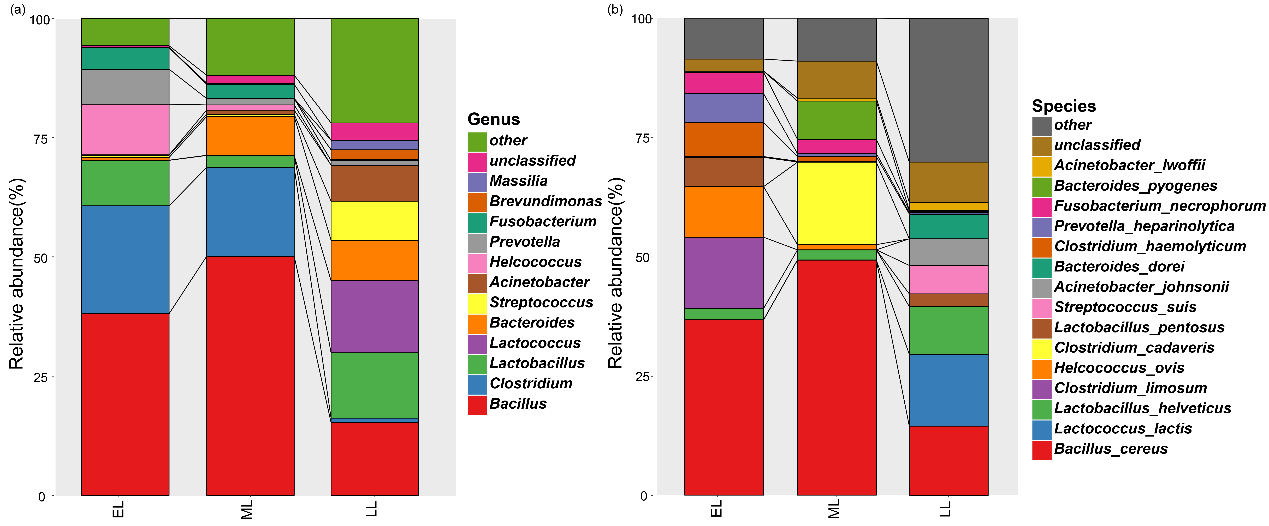

**Figure S2.** Composition of milk bacterial microbiota at the genus (A) and species (B) levels. Early, middle, and late lactation are represented by ‘EL’, ‘ML’, and ‘LL’, respectively.
